# PepMSND: Integrating Multi-level Feature Engineering and Comprehensive Databases to Enhance in vivo/in vitro Peptide Blood Stability Prediction

**DOI:** 10.1101/2024.12.12.628290

**Authors:** Hu Haomeng, Chengyun Zhang, Xu Zhenyu, Hongliang Duan

## Abstract

Deep learning technology has revolutionized the field of peptides, but key questions such as how to predict the blood stability of peptides remain. While such a task can be accomplished by experiments, it requires much time and cost. Here, to address this challenge, we collect extensive experimental data on peptide stability in blood from public databases and literature and construct a database of peptide blood stability that includes 635 samples. Based on this database, we develop a novel model called PepMSND, integrating KAN, Transformer, GAT and SE(3)-Transformer to make multi-level feature engineering to make peptide stability prediction. Our model can achieve the ACC of 0.8672 and the AUC of 0.9118 on average and outperforms the baseline models. This work can facilitate the development of novel peptides with strong stability, which is crucial for their therapeutic use in clinical applications.

## Introduction

Peptides and proteins gradually have become a popular modality in the pharmaceutical industry. Until now, there have been 80 approved peptide drugs applied to the clinic treatment. However, although these drugs have achieved remarkable achievements in clinical practice, no significant increase in the number of such drugs has been observed recently.^1^ This can be attributed to the instability of peptides. They are very easy to be hydrolyzed by proteases in the body, hindering the application of candidates in the clinical application.^2^ The degradation of peptides not only primarily occurs in the plasma, but also in the gastrointestinal tract, liver, and immune cells. This usually results in a very short half-life, making oral application unaccessible.^3–5^ To improve their stability, many modification strategies have been proposed: D-form or unnatural amino acid residues, N-methylation or forming cyclic peptides and conjugating with macromolecular carriers like proteins, lipids, and polymers.^6–8^

Considering the importance of the blood stability of peptides in their clinic application, how to measure/predict this property becomes an appealing issue.^9^ Traditionally, experimental approaches such as blood stability and enzymatic degradation tests are universal way. These methods can identify and evaluate the blood stability of peptides in different experimental settings with high accuracy. However, They necessitate high costs and long time, which can’t satisfy the recommendation for high throughput screening or large scale study.^10^ To address such a problem, people turn to computational methods, which have attracted much attention in other fields. Take ProtParam as an example, this technique explores half-life and N-termi residues based on N-end rule,^11–14^ and combines with the experimental statistics-based rule^15^ to measure the peptide stability. Additionally, a multi variable regression model also can be implemented to predict the half-life of peptides in blood.^16^ With the advancement of deep learning in peptide development, Mathur et. al ^17^ predict this property of peptides by using an SVM model that trained on a database consisting of the half-life of 261 peptides in mammalian blood.

Nevertheless, peptide blood stability prediction still faces essential challenges despite related developments in the past decades. For example, in the blood stability test, SUPR peptide showed a very different half-life when facing different experimental conditions: this peptide can be hydrolyzed in mouse plasma but stabilize in human plasma within 24 hours. However, such peptide blood stability differences usually be neglected due to the individual and fragmented data availability, which may lead to the model’s misclassifications. Additionally, many methods prefer to adopt relatively simple low-dimensional representation to illustrate the peptide feature, which usually neglects their conformation that is vital for distinguishing the activity difference.^18,19^ Actually, linear and cyclic peptides share the same amino acid sequences but have entirely different blood stability.^20^ To address the problem above, comprehensive systematized experimental data on peptide blood stability is necessary, which can accelerate the development of related research.

Therefore, in this study, we collect experimental data from public databases and literature as much as possible to build a specific peptide blood stability database. Furthermore, we make a comprehensive peptide feature engineering including basic molecular descriptors, SMILES, molecular graphs and complex 3D conformations to illustrate the intrinsic characters of peptides, thereby developing a novel model called PepMSND tailored for predicting peptide stability in various blood environments. The combination of our database and multimodal model offers an opportunity to identify potential peptide candidates with strong blood stability, improving the peptide drug development.

## Method

### Data collection

In this study, as shown in Figure 1, we collect peptide stability data samples manually from various sources such as published works, patents, and related databases. Based on the universal claim of Cavaco et.al that experimental half-life value is a good choice to demonstrate the stability of peptides,^21^ we adopt peptide[Title/Abstract] AND half-life[Title/Abstract] as the keyword to search associated information the PubMed and find 1413 works published in the range 2015-2024 year. In addition, we search public databases like PEPlife^22^, DrugBank^23^ and THPdb^1^ to explore more data. To ensure the quality and quantity of data, we make the following data clearing: 1. Removing peptides that lack or are missing stability information. 2. Removing peptides for which no explicit sequence information is given. 3. Excluding peptides for which experimental conditions are not explicitly given. 4. Ignorig peptides that are not experimented on in human or murine blood. 5. Excluding peptides with complex modifications (e.g., polyethylene glycol modifications). 6. Excludig peptides that cannot be represented by SMILES. Finally, a total of 635 samples are collected. Then, these peptides are classified into two classes: stable and unstable based on whether the peptides retained more than 50% of their intact fragments after 1 h in blood. In addition, we divide these samples into training and test datasets with a ratio of 9:1 and apply the 10-fold cross-validation to avoid potential bias.

**Figure 1.**
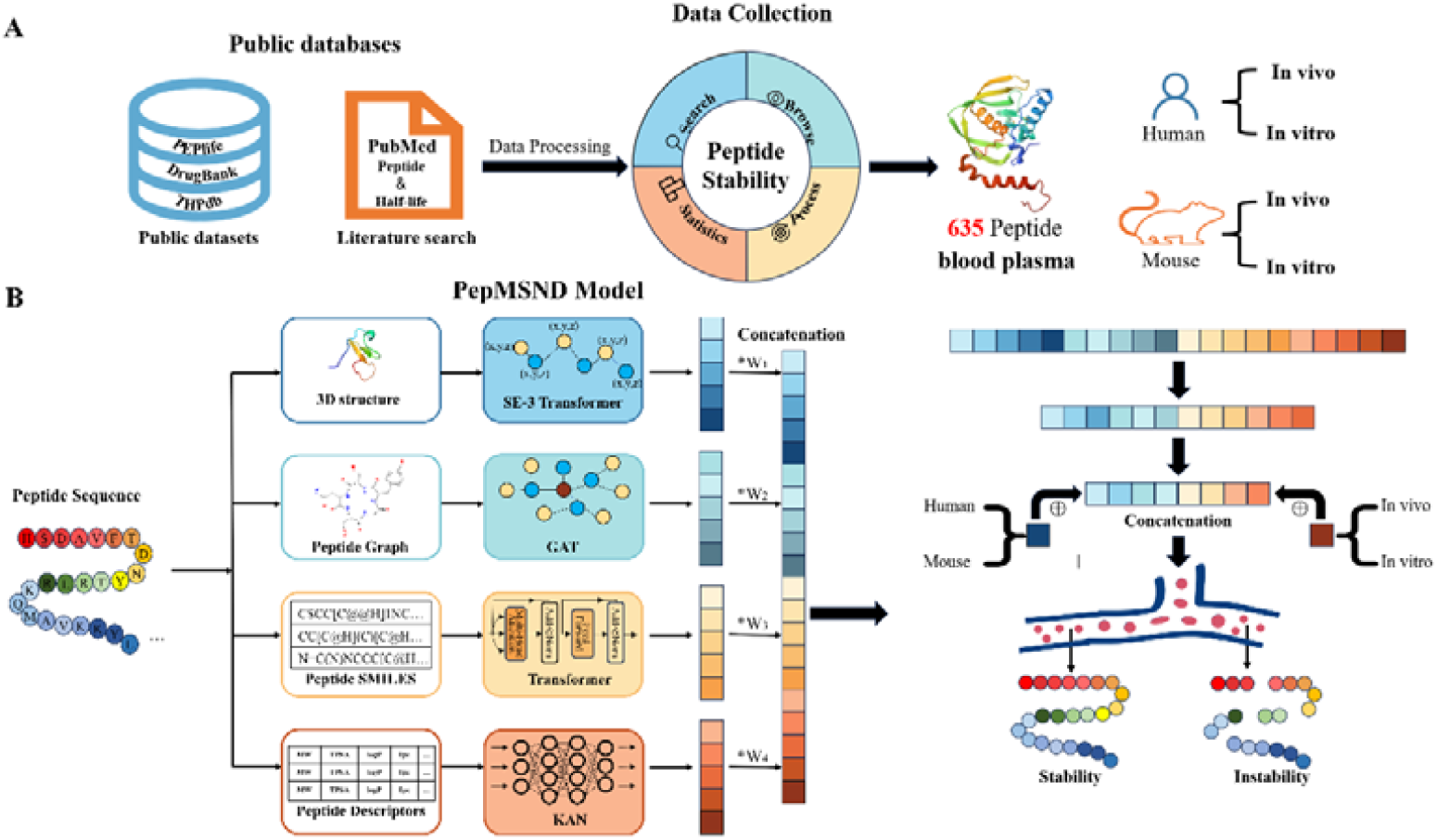
The workflow of our database and PepMSND model. (A) The illustration of the building process of the database. (B) The architecture of the PepMSND model. This model includes four modules: the SE(3)-transformer for the 3D peptide structure feature, the GAT model for the 2D peptide graph feature, the Transformer for 1D peptide SMILES and the KAN model for 0D peptide physicochemical properties.

FASTA format can’t afford the representation of peptides involving unnatural residues, N-methylation modification et.al. As displayed in Figure 2B, we use the

**Figure 2.**
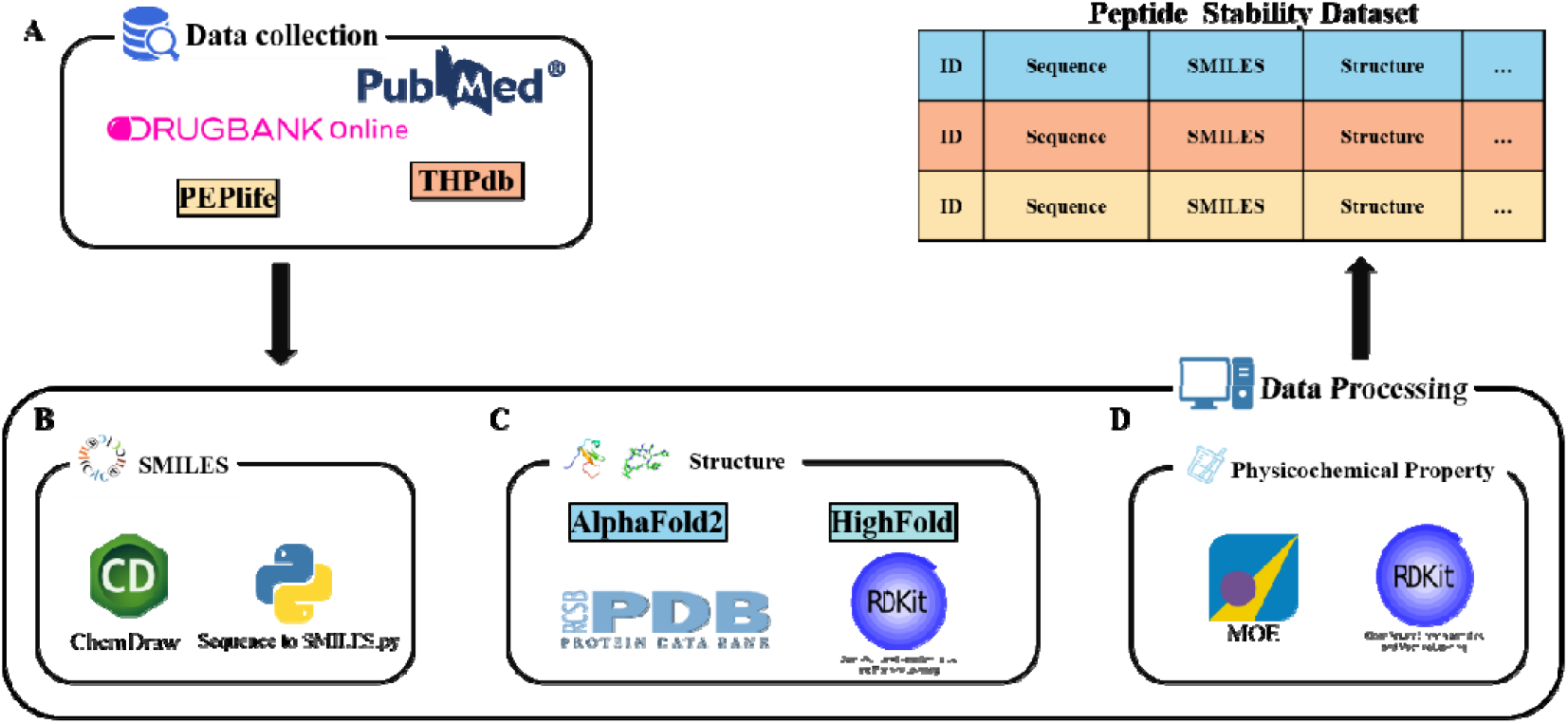
The illustration for the database building can be divided into two steps: data collection and preparation. (A) Data collection. (B) SMILES generation. (C) 3D structure generation. (D) physicochemical property calculation.

**Figure 3.**
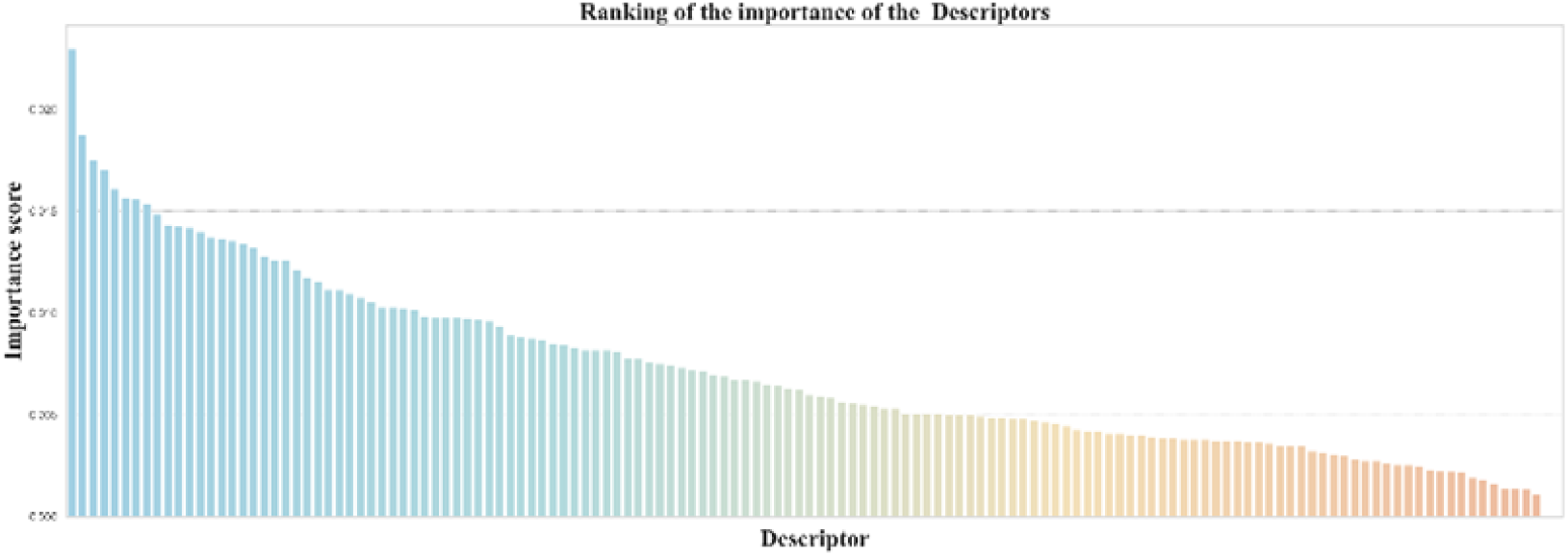
The top 140 peptide properties selected by RF.

SMILES to display the peptide feature at the atom level and propose a program to automatically convert FASTA sequences to SMILES. For those particularly complex structures, we choose to manually convert them using ChemDraw. All SMILES are standardized.

### Peptide structure generation

Traditionally, experimental observations like X-ray crystal diffraction, nuclear magnetic resonance spectroscopy, and electron microscopy are effective ways to investigate peptide structures.^24^ With the advancement of technology, structure prediction models like Alphafold^25^, RoseTTAFold^26^, ESMFold^27^ and HighFold^28^ can also provide plausible structures with high accuracy and efficiency. As displayed in Figure 2C, first, we search the PDB database, and for peptides that could not be retrieved, we adopt different strategies to deal with them. For the natural linear peptides, AlphaFold is implemented. As for the natural cyclic peptide, we use our proposed model HighFold. As for the peptides with complex modification, the RDkit (version 2023.3.2) is used. Based on this toolkit, 5000 conformations are generated for the peptide input and are optimized by the UFF force field. Ultimately, only the conformations with the lowest energy are selected for further experiments.

### The PepMSND model

Multimodal technology is a method that can efficiently integrate and process various data. It not only enhances the depth and breadth of data processing but also significantly improves the accuracy and generality of the model.^29^ In this study, we apply this technology to make a comprehensive feature engineering that takes physicochemical property, sequence information, molecular structure, and 3D conformation into consideration(Figure 1B). Specifically, molecular descriptors input as the 0D feature is processed by the Kolmogorov-Arnold Networks (KAN)^30^ to capture the physical and chemical properties of peptides;2. SMILES input as the 1D feature is processed by the Transformer^31^ to absorb the sequence relationship 3. molecular graph as the 2D feature is processed by GAT^32^ to learn the atoms and bonds interaction ; 4. Predicted structure as the 3D feature is processed by SE(3)-Transformer^33^ to provide additional information. Then, a series of learnable weights are used to integrate these features, thereby generating a joint feature vector that is followed by the further process in a shared layer. It should be noted that we introduce the species and experimental environment information to the model through one-hot coding, to pay more attention to these features. Detailed information about these models can be found in the following.

### KAN model

KAN (Kolmogorov-Arnold Networks) is proposed as a promising alternative to Multi-Layer Perceptrons (MLPs), which can learn both the compositional structure and the univariate functions quite well. This model adopts a learnable function parametrized as a spline to focus on edges (‘weight’), thereby improving the flexibility of models and allowing complex functions to be simulated with fewer parameters. In most study cases, smaller KAN can achieve comparable or better performance compared to MLPs. In our study, the KAN model is implemented to learn the peptide information about physicochemical properties presented by molecular descriptors. Specifically, we use the RDkit(version 2023.3.2) to calculate the associated property values and remove the descriptors including constant values to avoid information redundancy. Then, these selected descriptors are evaluated by a random forest model, and the top 140 important features are retained as the input for the KAN model. Additionally, we introduce two features: species and experimental environment. By utilizing one-hot encoding, these features are incorporated as additional 0D features into the set of molecular descriptors, thereby enriching the input information for the model. The function of this technology as follows:

### Transformer Model

Compared to molecule SMILES, peptide SMILES is longer and more complex. To effectively capture the intrinsic correlation with such a sequence, we adopt the Transformer. This model consists of three key layers: the embedding layer, attention layers and feed-forward neural network (FNN). In the embering layer, a positional encoder is introduced for precisely fusing the position information of each character in its sequence into the corresponding embedding vector. The function of this technology as follows: d_model_

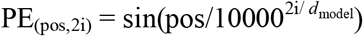

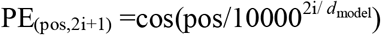

where, pos is position, i is Embedding vector dimension, *d*_model_ is model dimension.

The attention mechanism is the core of Transformerer, it can focus on the significant information among a vast amount of data and mitigate the impact of redundant information. Its calculation is shown in the following:

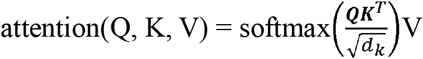

Where Q is the query, K is the key and V is the value, all are calculated based on inputted sequences.

### GAT Model

Graph Attention Networks (GATs) are neural networks specifically designed to handle graph data. During the message-passing process. GATs leverage the attention mechanism to dynamically learn the importance of neighbor nodes for each node This allows the network to pay attention to the most relevant information while ignoring less important or redundant information. In our task, the atoms are represented by nodes and bonds are represented by edges. With the implementation of GAT, the 2D feature can be absorbed. With its inherent ability to integrate graph structure information, GAT can comprehensively and deeply analyze the interrelationships and patterns among the atoms in the polypeptide. This feature enables us to extract meaningful features that provide strong support for subsequent classification tasks. Its calculation is shown below:

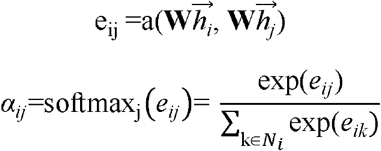

Where **W** is a weight matrix. 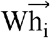 and 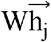 are the feature vectors of nodes i and j after linear translation. Ni is the neighbourhood set of node i, α_ij_ is the attention coefficient normalized by the softmax function.

### SE(3)-Transformer Model

The SE(3)-Transformer is a self-attention module variant for 3D point clouds and graphs. Under continuous 3D rototranslations, this model can remain equivariant. In the model, each atom is represented by its 3D coordinates (x, y, z). This model consists of three components. These are (1) edge-wise attention weight αij that SE(3) on each edge is unchanged, (2) edge-wise SE(3)-equivariant value messages which propagate information between nodes, (3) a linear/attentive self-interaction layer. The Attention for each structural node is calculated as follows:

Its calculation is shown below:

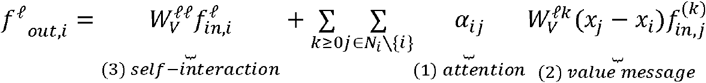

Where 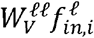 is the self-interaction term, showing the result of the input feature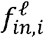 of node i after undergoing a linear transformation. αij is the attention weight, representing the degree to which node i pays attention to or attends to its neighboring node j. 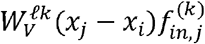 is the value message term, including the message sent from node j to node i. This includes the positional difference between nodes (x_j_−x_i_) and the input feature 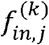 of node j.

It should be noted that a simplified version of the SE(3)-Transformer model that contains equilateral layers and 3D coordinates is adopted in our study to deal with the peptide structure.

### Baseline models

6 baseline models are chosen to compare to the PepMSND model.

Support Vector Machines (SVMs): SVM^34,35^ is a classic model that distinguishes different data points by finding the optimal hyperplane in the feature space.

Random Forest(RF): RF^35^ is one of the ensemble learning methods that guarantee its accuracy and robustness based on the overall consideration of multiple decision trees.

Extreme Gradient Boosting (XGBoost): XGBoost^36^ is a machine learning algorithm based on a gradient lifting framework, which aims to achieve efficient, flexible and portable distributed gradient enhancement.

K-Nearest Neighbors(KNNs): KNN^37^ is generally implemented to the data mining and image classification tasks. This model classifies data points by majority voting based on the nearest neighbors in the feature space.

Graph Isomorphism Network(GIN): GIN^38^ is an effective model for dealing with graph data. Its core advantage lies in its strong expression ability and deep understanding of heterogeneous graphs.

## Metric

In this study, we adopt several metrics: Accuracy (ACC), Precision(Pre), F1 score(F1_Score), Recall, Area Under the Curve(AUC), Matthews correlation coefficient(MCC) to evaluate the performance of models in predicting the blood stability of peptides:

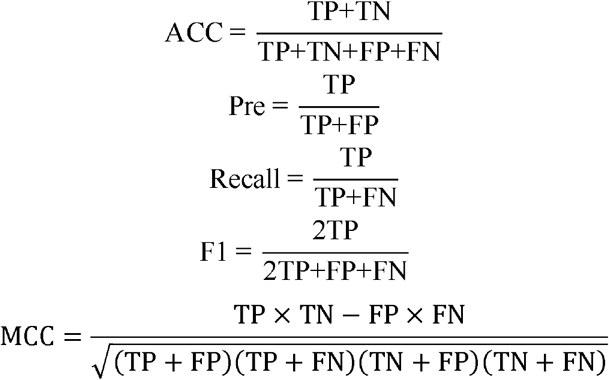

The numerical ranges for ACC, Pre, Recall, F1-Score, and AUC are all from 0 to 1, with higher values indicating better model performance. The numerical range for MCC is from -1 to 1, and similarly, higher values also represent better model performance.

## Results and Discussion

### Dataset

In our dataset, the number of cyclic peptides is 107 and that of linear peptides is 528(Figure 4C). Among cyclic peptides, many of them are cyclized by the disulfide bonds. To be more stable, most peptides have been modified in different ways including D-residue replacement or introduction of N-terminal capping. In addition, more complex modifications such as N-methylation also can be found in this dataset. Figure 5A displays that the lengths of most peptides range from 5 to 50 amino acids, accounting for 73% of the total dataset (Figure 4A). As shown in Figure 4D, in this dataset, the number of peptides evaluated in in vivo human blood is 115, while the number evaluated in in vitro human blood is 222. Additionally, the number of peptides tested in in vivo mouse blood is 217, and the number evaluated in in vitro mouse blood is 81, showing the diversity of our data. To more clearly illustrate the peptide stability distribution, we divide peptides into four categories: unstable, stable, highly stable, and non-degradable. Specifically, the stability of peptides can be classified according to the following criteria: a peptide is considered unstable if the proportion of the original peptide in the blood drops to less than 50% after 1 hour; if at least 50% of the original peptide remains unhydrolysed from 1 to 6 hours, the peptide is classified as a stable peptide; if the proportion remains above 50% from 6 to 12 hours, it is regarded as highly stable; and if the original peptide remains unhydrolysed for more than 12 hours, it is classified as a non-degradable peptide. As shown in Figure 4B, the majority of peptides (approximately 57% of the dataset) belong to the unstable class. As for the stable peptides, most of them have some modification such as unnatural residue substitution.

**Figure 4.**
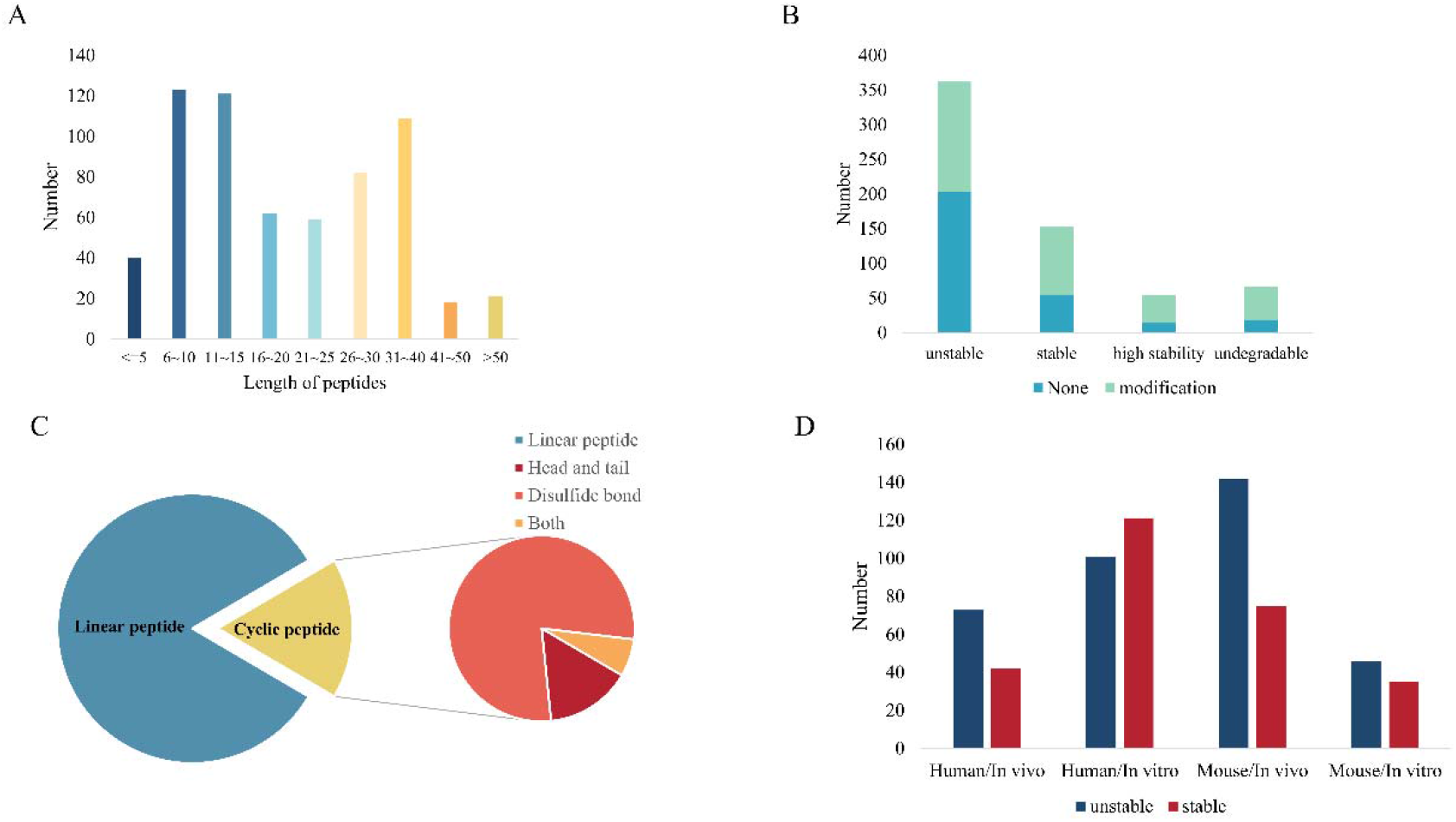
The distribution of the database. (A)The length distribution of peptides. (B)The distribution of natural and unnatural peptides. (C)The distribution of cyclic and linear peptides. (D)The distribution of peptides in different blood environments.

**Figure 5.**
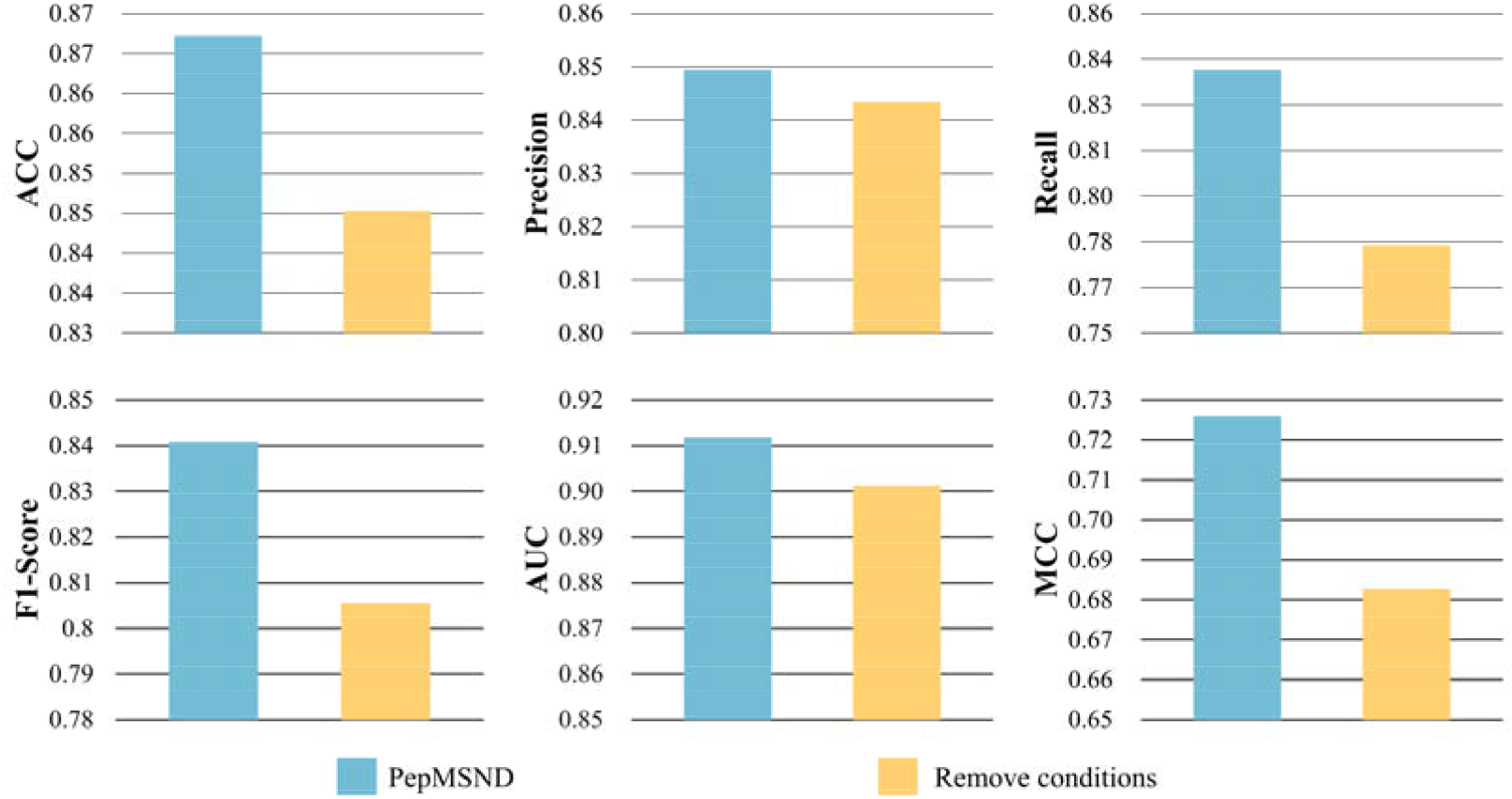
The performance comparison before and after removing blood experimental environment information.

**Figure 6.**
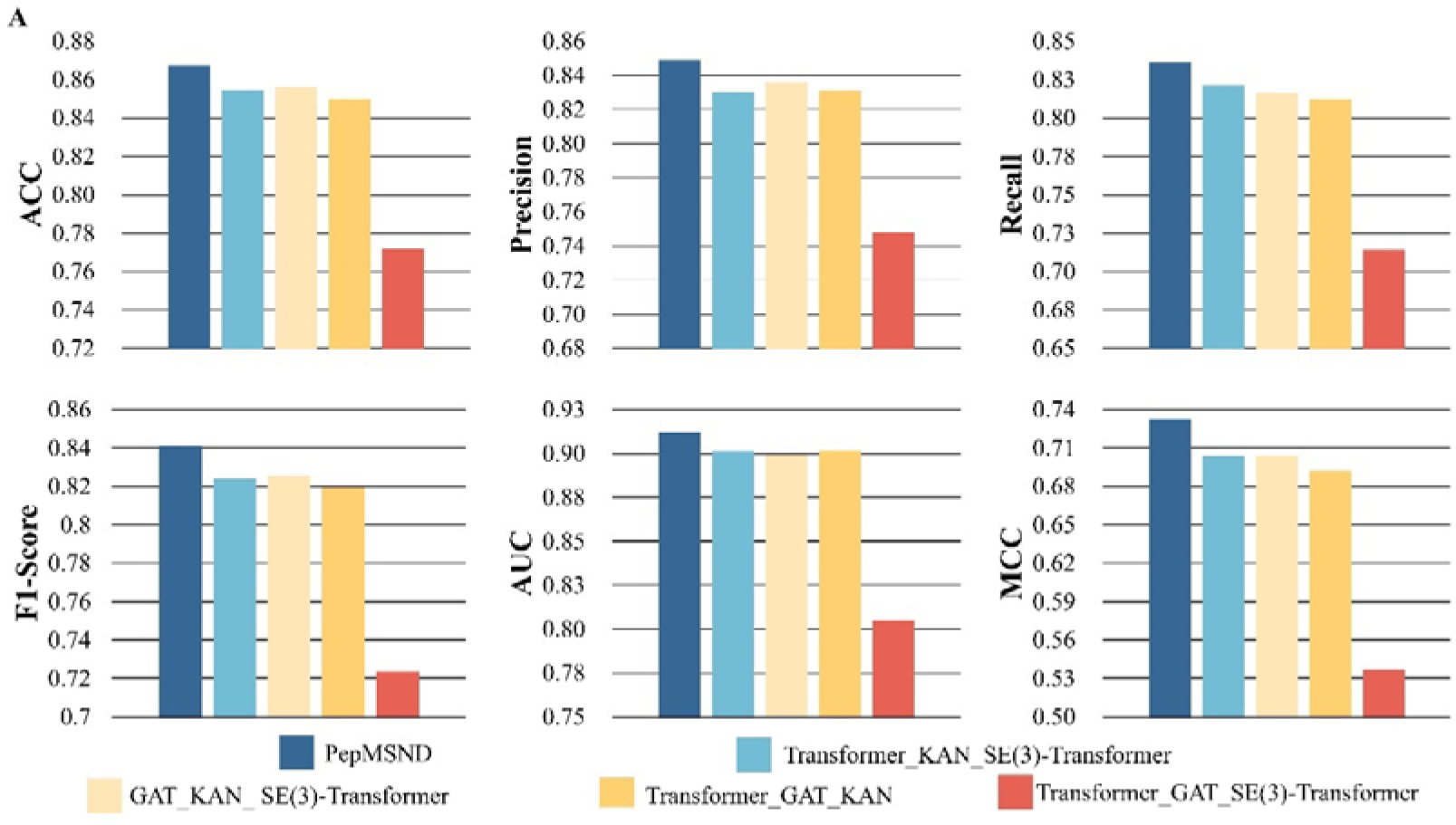
Results of the ablation experiment.

### Performance comparison

Based on this dataset, we develop a novel model called PepMSND to predict blood stability. To comprehensively evaluate the performance of this model, we compare it to several different models. In this comparison, all models are trained on the same datasets and evaluated by the 10-cross validation. Table 1 displays the results of models in this task.

**Table 1.**
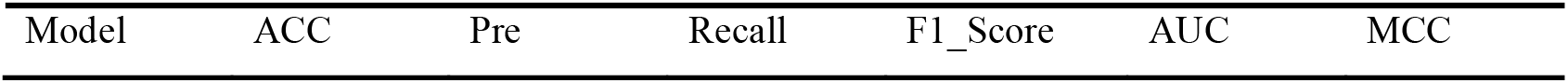
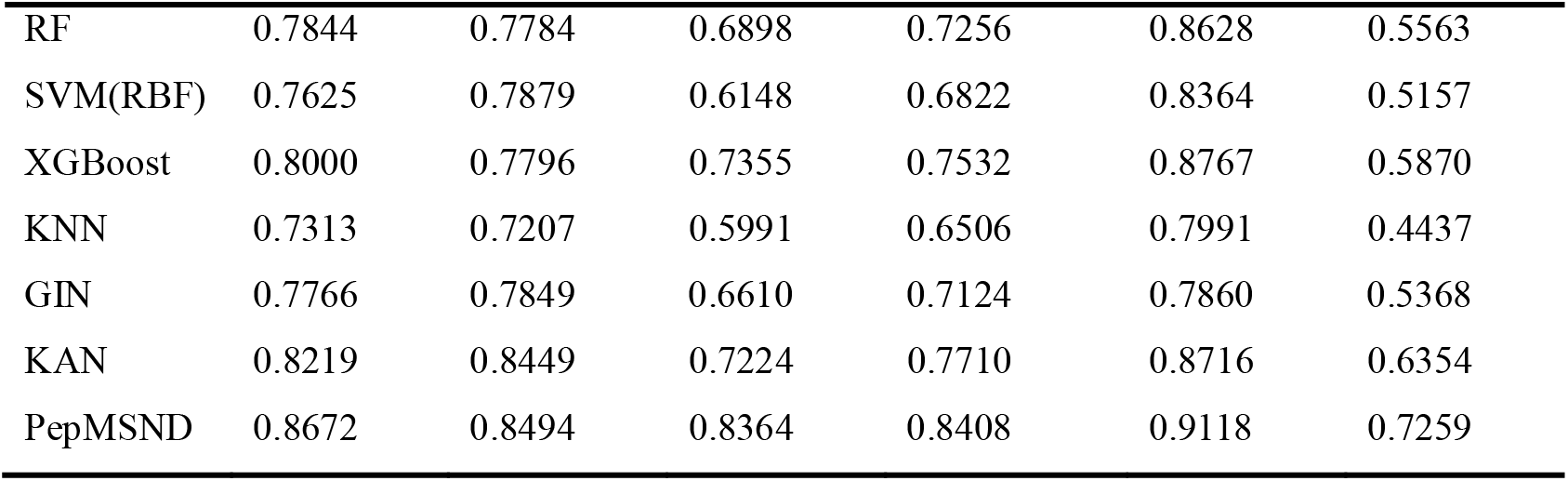
Performances of various models with different metrics on the test set. These results are all average values of 10-fold cross-validation.

**Table 2.** Performances of different species and experimental environments with different metrics on the test set.

|  | ACC | Pre | Recall | F1_Score | AUC | MCC |
| --- | --- | --- | --- | --- | --- | --- |
| Human/In Vivo | 0.9186 | 0.8940 | 0.8821 | 0.8671 | 0.9045 | 0.8269 |
| Human/In Vitro | 0.8267 | 0.8578 | 0.7825 | 0.8109 | 0.8669 | 0.6476 |
| Mouse/ In Vivo | 0.8883 | 0.8352 | 0.8634 | 0.8457 | 0.9298 | 0.7567 |
| Mouse/ In Vitro | 0.8937 | 0.8750 | 0.9357 | 0.9005 | 0.9458 | 0.7971 |
| PepMSND | 0.8672 | 0.8494 | 0.8364 | 0.8408 | 0.9118 | 0.7259 |

These results are all average values of 10-fold cross-validation.

Table 1 demonstrates that PepMSND exhibits excellent performance in these metrics, showing 0.8672, 0.8494, 0.8364, 0.8408, 0.9118, 0.7259 in the ACC, Pre,

Recall, F1_Score, AUC, and MCC, respectively. Compared to other models, PepMSND demonstrates superiority in all metrics. Especially in AUC, PepMSND reaches 0.9118, which demonstrates its ability to distinguish positive and negative samples when faced with unbalanced datasets. In addition, we observe that the traditional descriptors-based models often perform better than graph-based models. It reveals that the peptide physicochemical properties are very important for this task and more training samples are required for the graph-based models to capture associated information.

### The effect of experimental environments on model performance

We are also interested in the performance of our model in different experimental blood environments. Based on different species and experimental environments, we divide our test dataset into four classes. In most cases, PepMSND demonstrates satisfactory ability in predicting the stability of peptides with metric values higher than 0.8. Take the prediction in the in vivo human blood condition as an example, the PepMSND achieves 0.9186, 0.8940, 0.8821, 0.8671, 0.9045, 0.8269 in the ACC, Pre, Recall, F1_score,

AUC and MCC, which display the great power of this model to understand the peptide in this environment. The stable model performance indicates that our model delivers high accuracy with great generality. But we also notice that the change in the experimental environments also affects the model. The ACC gaps between Human/In Vivo and Human/In Vitro can reach 9.19%. However, such a phenomenon is not investigated in the experiments with Mouse/In Vivo and Mouse/In Vitro. Additionally, the species also is one factor affecting the model’s ability. Compared to the ACC of Mouse/In Vitro environment, the ACC of Human/In Vitro is lower.

In previous studies, most researchers often ignored the potential impact of species and experimental environment differences on the model. While many stabilities similarities can be investigated across species and in vivo and in vitro blood stability experiments for the same peptides, some peptides show very different stability results in different experiment conditions, leading to missing potential compounds. Therefore, we emphasized the importance of the experimental environment to the model during the model building. To demonstrate the importance of these two pieces of information, we explore the model performance when removing them. The results are displayed in Figure 5. After the removal of these two conditions, significant drops appear in all evaluation indicators. The decrease in ACC, Recall, F1_Score, AUC, and MCC is 2.19%, 5.76%, 3.54%, 2.38%, and 4.32%, respectively, showing the great importance of blood experimental environment information.

### The effect of peptide length on model performance

We are also interested in the influence of peptide length on the model performance. Therefore, in this section, we make a deep analysis of the model performance in the peptides with different lengths. As shown in Table 3, the PepMSND model demonstrates its generality in these different subsets, achieving all ACC exceeding 0.8. Especially in predicting the stability of peptides with lengths of 25-40, the ACC of our model can reach 0.9352, demonstrating great predictive ability. In other metrics, this model also gains satisfactory results: 0.9300 of Pre, 0.8603 of Recall, 0.8803 of F1_Score, 0.9696 of AUC and 0.8445 of MCC. This may be attributed to the training dataset that Most of the peptides contained are in length of 25-40. Therefore, compared to other peptides, the PepMSND model has a deeper understanding of peptides with 25-40 lengths.

**Table 3.** Performances of different lengths with different metrics on the test set.

|  | ACC | Pre | Recall | F1_Score | AUC | MCC |
| --- | --- | --- | --- | --- | --- | --- |
| 5~25 | 0.8179 | 0.8116 | 0.8076 | 0.8055 | 0.8648 | 0.6376 |
| 26~40 | 0.9352 | 0.9300 | 0.8603 | 0.8803 | 0.9696 | 0.8445 |
| 5~40 | 0.8571 | 0.8379 | 0.8314 | 0.8329 | 0.9047 | 0.7063 |
| PepMSND | 0.8672 | 0.8494 | 0.8364 | 0.8408 | 0.9118 | 0.7259 |

These results are all average values of 10-fold cross-validation.

### Ablation Experiment

In this study, we use different modules to focus on different peptide features. Here, we conducted an ablation study to explore the effect of these different modules. Under the same experimental setup, we implement several variants of the PepMSND model for predicting peptide stability. These include the following:

(1) Transformer_KAN_SE(3)-Transformer: only containing Transformer, KAN and SE-3Transformer,
(2) GAT_KAN_SE(3)-Transformer: only containing GAT, KAN and SE(3)-Transformer,
(3) Transformer_GAT_KAN: only containing Transformer, GAT and KAN,
(4) Transformer_GAT_SE(3)-Transformer: only containing Transformer, GAT and SE(3)-Transformer.

Figure 5 shows the performance comparisons of PepMSND and its variants. No surprise, the lack of one module can lead to a decline in PepMSND’s performance. Compared to the Transformer_KAN_SE(3)-Transformer, GAT_KAN_SE(3)-Transformer, Transformer_GAT_KAN, Transformer_GAT_SE(3)-Transformer, our model displays a deeper understanding of this tasks, achieving the F1 improvement of 1.70%, 1.52%, 2.1%, 11.72%, respectively. Such improvement is also can be observed in the ACC metric, where our model reaches increases of 1.25%, 1.1%, 1.72%, 9.53%. Furthermore, we make a deep analysis to explore the effect of these modules in our model. As demonstrated in the table, both the Transformerer_KAN_ SE(3)-Transformer and the GAT_KAN_ SE(3)-Transformer exhibit comparable performance in predicting the stability of peptides. This indicates that both the Transformers and GAT architectures play equally significant roles in our model. It should be noted that Transformer focuses on the sequence feature, which is rather different from the graph-based GAT. The former contributes to finding the correlation between tokens in a sequence and the latter captures the explicit information about atoms and bonds. Namely, the text feature (1D feature) is as same as the graph feature (2D feature). We also noted that the Transformer_GAT_KAN that not contain the

SE(3)-Transformer module which lacks the SE(3)-Transformer module, performs inferior to the aforementioned two models, achieving an ACC of 0.8500, F1 score of 0.8198. This underscores the significance of the 3D structure of peptides in our model. When the KAN module is removed from our model, we observe a more significant performance decline across all metrics, with a decrease of more than 0.1. This decrease is most pronounced in the MCC metric, which drops from 0.7259 to 0.5370. This suggests that the model without the KAN module has a worse ability to predict peptide blood stability. The inclusion of the KAN module appears to be crucial for the model’s performance, likely due to its ability to capture important features and relationships involving different physicochemical properties. In other words, these peptide properties can assist the model in establishing a correlation between peptides and their stability. By incorporating these properties into the model, our model can better understand and predict how different peptides will behave in the blood, which is important for various applications such as drug design and peptide-based therapies.

## Conclusion

In this study, we provide a comprehensive peptide blood stability database. This dataset has diverse blood experiments and peptides, comprising 635 samples. It represents the first open-source database that offers easy access to peptide stability in blood. Furthermore, we develop a multi-model model based on this dataset to predict peptide blood stability across various experimental settings. This model integrates multi-dimensional features, including 0D physicochemical property, 1D sequence, 2D graph, and 3D structure, to capture the correlation between peptides and their stability.

To evaluate the ability of our model, we compare it to baseline models such alike GIN. In this performance comparison, our model exhibits an excellent ability to determine whether the peptide is stable in blood and achieves 0.8672, 0.8494, 0.8364, 0.8408, 0.9118, 0.7259 in ACC, Pre, Recall, F1_Score, AUC, MCC respectively. Additionally, our model shows improvements of 4.53% in ACC, 6.97% in F1_Score, 4.02% in AUC, and 9.05% in MCC, when compared to the best baseline model. This benchmark experiment demonstrates our model’s superiority in predicting peptide stability. To further explore the effect of different features in our model, we make a series of ablation experiments. The results show that in this model, 1D sequence information and 2D structural information play equally important roles, while 3D structural information and 0D physicochemical property information play a more important role and can provide more useful information. However, although these variant models perform worse than our model, they still show superiority when compared to other baseline models such as SVM.

It is worth noting that the peptide stability dataset exhibits inferior quality and quantity compared to the molecule stability dataset, thereby posing significant obstacles to the practical deployment of our model in real-world scenarios. Specifically, our peptide stability database necessitates continuous updates. Furthermore, the implementation of a transfer learning strategy may represent an effective solution to alleviate such data scarcity challenges, considering the power of this method in other fields. Additionally, due to the current limitations of the structure prediction model, accessing a high-precision peptide structure model remains challenging.

## Code availability

The code and dataset will be released at GitHub after publication.

